# Performance of an Ambulatory DRY-EEG Device for Auditory Closed-Loop Stimulations in the Home Environment

**DOI:** 10.1101/181529

**Authors:** E. Debellemaniere, S. Chambon, C. Pinaud, V. Thorey, D. Léger, M. Chennaoui, P.J. Arnal, M.N. Galtier

## Abstract

**Objective:** Recent research has shown that auditory closed-loop stimulations can enhance sleep slow oscillations (SO) to improve N3 sleep quality and cognition. Previous studies have been conducted in a lab environment and on a small sample size. The present study aimed at validating and assessing the performance of a novel ambulatory wireless dry-EEG device (WDD), for auditory closed-loop stimulations of SO during N3 sleep at home.

**Material and Methods:** The performance of the WDD to detect N3 sleep automatically and to send auditory closed-loop stimulations on SO were tested on 20 young healthy subjects who slept with both the WDD and a miniaturized polysomnography (part 1) in both stimulated and sham nights within a double blind, randomized and crossover design.

The electrophysiological effects of auditory closed-loop stimulation on delta power increase were assessed after 1 and 10 nights of stimulations on an observational pilot study in the home environment including 90 middle-aged subjects (part 2).

**Results:** The sensitivity and specificity of the WDD to automatically detect N3 sleep in real-time were 0.70 and 0.90, respectively. The stimulation accuracy of the SO ascending-phase targeting was 45±52°. The stimulation protocol induced an increase of 39.5 % of delta power after the stimulations. The increase of SO response to auditory stimulations remained at the same level after 10 consecutive nights.

**Conclusion:** The WDD shows good performances to automatically detect in real-time N3 sleep and to send auditory closed-loop stimulations on SO accurately. These stimulations increased the SO amplitude during N3 sleep without any adaptation effect after 10 consecutive nights. This tool provides new perspectives to figure out novel sleep EEG biomarkers in longitudinal studies and can be interesting to conduct broad studies on the effects of auditory stimulations during sleep.

## I. INTRODUCTION

Sleep is a complex process that plays a key role in maintaining homeostasis, well-being and overall health [1]–[3]. In recent decades, increasing evidence has confirmed that slow-wave sleep (SWS) had a major impact in many biological functions such as glucose metabolism, hormone release, immunity and memory [4]–[7]. This proposed role for SWS, coupled with observations of impaired SWS in several chronic pathologies such as fibromyalgia [8], as well as in aging [6], [9] have led to imagine the development of methods that could specifically enhance SWS (see [10] for Review). Several treatments, in which recently Tiabagine have been tried to increase slow oscillations (SO) of SWS [11], [12]. Transcranial direct current stimulation and transcranial magnetic stimulation have also been shown to be able to induce slow waves [13]. However, given the impractical and hazardous aspects of these methods, especially for chronic long-term exposure, more attention has been given to the possibility of enhancing slow waves by using more physiological stimuli. Among different sensory modalities, vestibular stimulations [14] and auditory stimulations appeared to be effective in increasing the magnitude of SO [15]–[21].

However, given the few possibilities to analyze electroencephalography (EEG) sleep in real-time and the constraint imposed by closed-loop stimulations, these studies have been generally conducted in small groups of subjects, only in a laboratory environment and with only few nights of polysomnography (PSG). As raised by a recent study, this standard practice that involves EEG monitoring in appropriate sleep infrastructures requires an important monetary, time and trained human resources costs for the development of the stimulation algorithm, the EEG hook-up, the overnight supervision, the triggering of the stimulation algorithm through the night, the EEG unhook-up and the sleep scoring [22].

While numerous EEG devices have engaged in developing lifestyle EEG solutions [22], fewer EEG devices have been specifically developed for sleep purposes trying to both record EEG recording and to automatically sleep score [23]. Among them, the now-discontinued Zeo device (Zeo, Inc., Newton, MA) seemed to be one of the most effective devices with scientific performance assessment. The studies evaluating its performance concluded that the Zeo device was useful for sleep monitoring at home with some weaknesses related to the over scoring of REM sleep and the underestimation of wakefulness [24], [25], [26], [27], [9], [28]. To our knowledge, there are no available tools on the market to analyze sleep EEG in real-time which can send auditory closed-loop stimulations on SO.

The aims of our study were to assess i) the performance of the Beta version of the Wireless Dreem Device (WDD) to detect N3 sleep automatically and to send auditory closed-loop stimulations on SO as compared to gold standard miniaturized polysomnography (PSG) (part 1) and ii) to test the electrophysiological effects of auditory closed-loop stimulation on a larger cohort in an observational pilot study at home (part 2).

## II. MATERIALS AND METHODS

### A. Subjects and settings

#### - Part 1: Validation of the acquisition and accuracy performances of the WDD in a clinical study

Twenty-four healthy subjects were recruited and included in this clinical trial through local and universities advertisements. This experiment was performed by the Alertness, Fatigue and Sleep Team (EA 7330) in the Hôtel Dieu Hospital. The local ethics committee approved the experimental protocol and complied with the tenets of the Declaration of Helsinki (Number of clinical trial: NCT02956161). All volunteers gave their informed written consent prior to participation. They received a monetary compensation for their time. Routine surveys and a medical interview with a physician ensured that they were non-smokers, had no history of neurological, psychiatric or endocrine disease, including any sleep disorder. All participants were free from medication except hormonal contraceptives. They were asked to follow a regular sleep/wake rhythm for at least 4 weeks prior to the experiment with 7–10h per night and no daytime naps. Their sleep and wake patterns were assessed with a sleep agenda and a wrist-actimeter (Actiwatch TM; Cambridge Neurotechnology, Cambridge, UK) from one-week prior the beginning of the experiment to the end of the protocol.

Subjects underwent 4 ambulatory home night’s monitoring with both the WDD and a PSG. The PSG was set at the sleep laboratory between 5 to 8pm. Participants were asked to put the headband and launch the recording themselves prior to sleep. In the morning, they were asked to remove the electrodes and bring the material back to the sleep lab.

The first night consisted in a habituation night that was discarded from the analyses. The three following nights encompassed: i) Sham condition where SO during N3 sleep was targeted but no sound was triggered ii) Ascending condition where the ascending phase of the SO during N3 sleep was targeted with sounds triggered and iii) Random condition where stimulations were played on the ascending, descending, up and down phases of SO were randomly targeted during N3 sleep. This study was double blind, randomized and crossover designed. The washout period was 1 week between each condition.

#### - Part 2: Electrophysiological effects of auditory closed loop stimulation provided by the WDD in an observational pilot study

The study consisted in an observational pilot study on subjects who bought and used the beta version of the Dreem headband ((Rythm sas, Paris, 2016), referred as the WDD in the manuscript) from November 2016 to June 2017. Since buying the device was a voluntary act, no exclusion criteria were followed except having a sleep or neurological disorder (assessed by a questionnaire). Similarly, the number of nights spent with the headband was not controlled and the choice to wear the headband was left to the subject.

To filter out the inherent bad recordings due to the home environment which is not as controlled as in laboratory, recordings with a minimum duration of 5h, a minimum effective sleep time of 3h and with a good EEG signal quality (higher than 60% of the time) were considered (see algorithm description below). To avoid the impact of outliers, recordings without N3 sleep or with more than 3h of N3 sleep were removed from the analysis.

### B. Materials

#### - Polysomnographic recordings

The PSG device consisted in miniaturized multi-channel ambulatory recording devices (Actiwave®, CamNtech Ltd England) with the following derivations: 6 EEG: Fp1-M2, C3-M2, O1-M2, Fp2-M1, C4-M1, O2-M1, 2 electro-oculograms (EOG), 2 chin electromyograms (EMG) and an electrocardiogram (ECG) [29]. The habitual frontal derivations were replaced by Fronto Polar position (FP) in order to be placed right next to the headband electrodes position. Bio-electrical signals were digitized at a sampling frequency of 128Hz with a 10-bit quantization between −500 and +500 μV, within a bandwidth of 0 to 48Hz. All the data were stored into computer files using the standard .EDF data format. EEG cup-electrodes (Ag-AgCl) were attached onto the scalp of the subjects (EC2 electrode cream, Grass Technologies, An Astro-Med, West Warwick, USA), according to the international 10-20 system for electrodes placement. Auto-adhesive electrodes (Neuroline 720, Ambu A/S, Ballerup, Denmark) were used for EOG recordings. The recording devices were fixed on the participant’s’ head using EC2.

#### - Ambulatory dry-EEG device: The WDD

The WDD device is a wireless system using 5 dry nanocarbon-coated fabric sensors to record EEG signal in ambulatory. The 4 EEG derivations were FPp1-M1, Fp2-M2, Fp1-Fpz and Fp1-Fp2 where Fpz was the virtual ground (Fig.1). The 2 derivations used for sleep analysis were FPp1-M1 and Fp2-M2. Unconventionally, the derivations were not contra-lateral wired because unilateral derivations improve the signal quality of the WDD by limiting electrodes disbond artifacts. Indeed, when sleeping on their side, it is generally one entire side that is artifacted. The WDD has a unique size with an elastic band behind the head that makes it adjustable such that it is tight enough to be secure, but loose enough to minimize discomfort. The signal is measured at 250Hz, filtered in the 0.4 to 18Hz band, and post processed according to the algorithms described below. An accelerometer is embedded in the WDD providing accurate positioning of the head at a sampling frequency of 50Hz. A bone conduction device integrated in the frontal band of the WDD on the forehead delivers sounds. This minimizes the sound that travels through air while keeping a perceptive sound loud enough for the headband wearer.

**Fig. 1.**
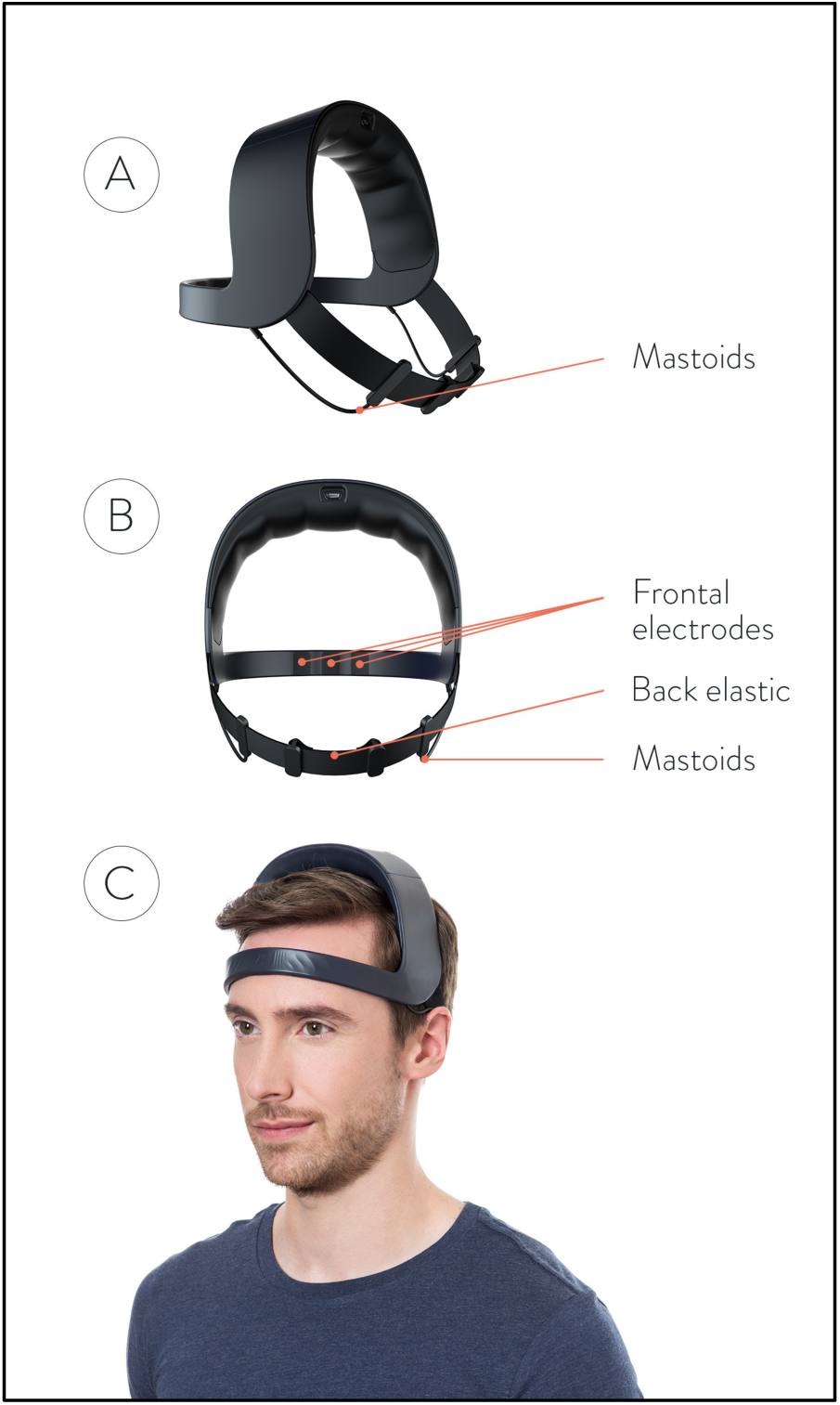
Representation of the WDD. A- Front view, B- Back view, C- Side view. The device is made up of 4 dry measuring electrodes: two front sensors placed in Fp1, Fp2 and two “reference electrodes” placed behind the ears as “mastoids” electrodes. The top arch gathers all the electronic components.

### C. Data analysis

#### - Embedded real-time algorithms

To produce auditory stimulations at a precise moment, the WDD implements a complex pipeline of operations, which is presented in a simplified form and detailed block by block below (Fig. 2A, B, C, D, E). Overall, the 3 inputs of the pipeline are the two frontal-mastoid EEG derivations x1 and x2 and the three-dimensional accelerometer variable denoted a. Both EEG channels are filtered a priori with a combination of infinite impulse response filters (Fig. 2A). More precisely, the signals were filtered with the following causal filters: a 4th order bandpass butterworth filter in the 0.4 to 4Hz frequency band, a 58 to 62Hz bandstop 6th order butterworth filter, a 48 to 52Hz bandstop 6th order butterworth filter and a 62 to 63Hz bandstop 2nd order Bessel filter.

**Fig. 2.**
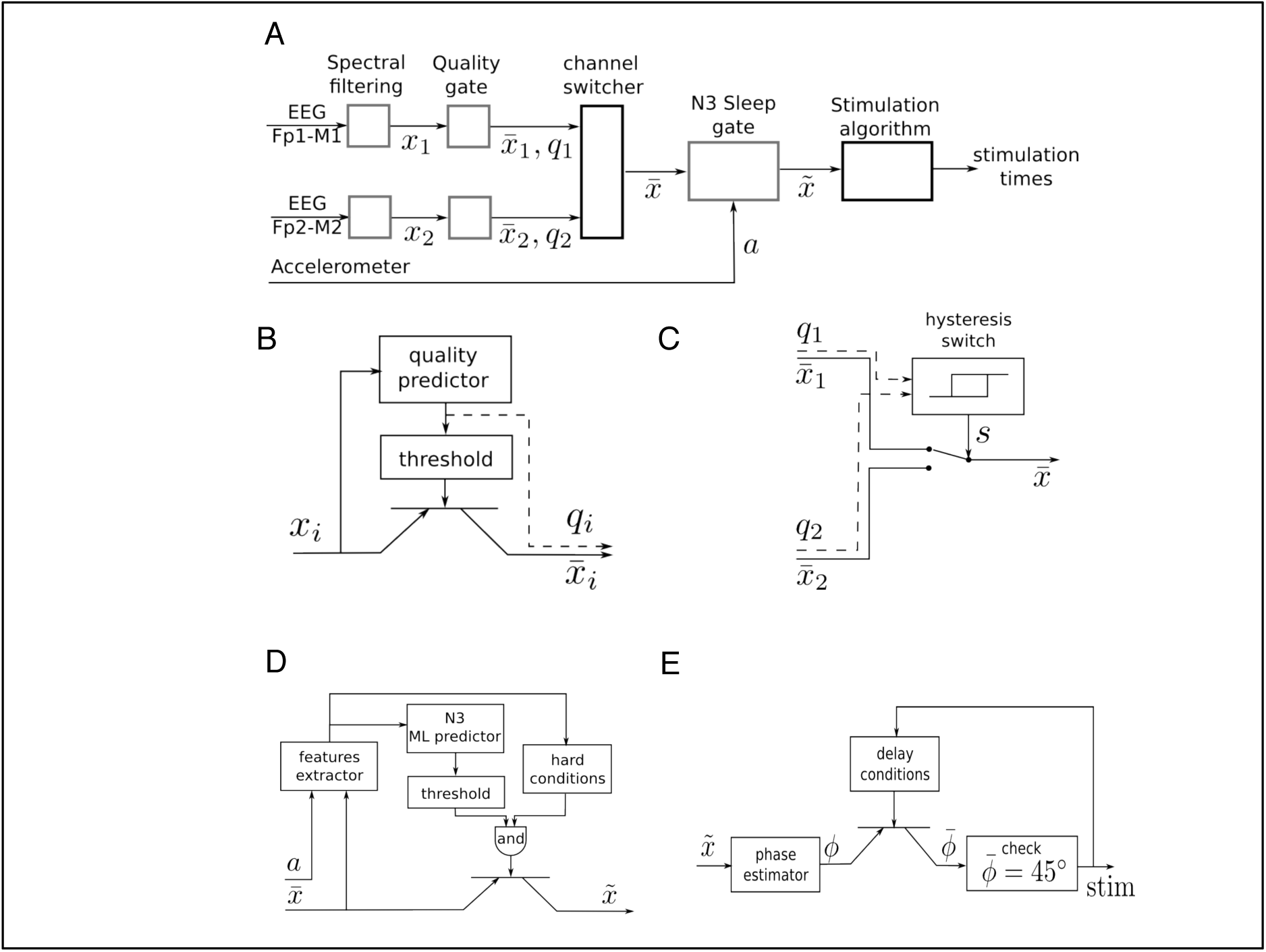
Coarse-grained algorithmic pipeline used by the WDD for stimulating N3 sleep. **A**- General pipeline. **B**- Quality gate representation, High-level block diagram of the quality gate. The filtered signal x_i_ (with i∈{1,2}) are sent to the quality predictor which computes an index of the signal quality q_i_∈[0,1] (0 means bad signal and 1 means perfect signal). This is a quantification of the extent to which the signal is perturbed by external artifacts, e.g. due to dry electrode bad contact. This quality index is the compared to a threshold to open or not the quality gate, which is illustratively represented via a transistor symbol. The output signal 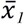 is equal to x_i_ if q_i_ > {threshold} and is Not a Number (NaN) else. **C-** High-level block diagram of the channel switcher. The two quality indices q_1_ and q_2_ enter an hysteris switch which selects which inputs, 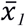 or 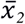, are broadcasted to the next block. The hysteris switch is parameterized by a threshold θ and outputs a binary variable at time t computed as s(t) = 1 if q_1_ - q_2_ > θ or (-θ < q_1_ - q_2_ < θ and s(t-1) = 1). Symmetrically, s(t) = 0 if q_1_ - q_2_ < -θ or (-θ< q_1_ - q_2_ < θ and s(t-1) = 0). If s(t) = 1 then 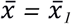 and, if s(t) = 0 then 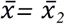. **D-** N3 Sleep gate representation. High-level block diagram of the N3 sleep gate. Both inputs, the accelerometer a∈R^3^and the virtual channel 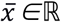, are used in a series of operations to identify whether, the input corresponds to N3 sleep, in which case the virtual channel is broadcasted to the next block. More precisely, a and 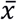 enters a series of signal processing functions in the feature extractor. Not only are basic statistics of the two signals computed, but also some classical patterns of N3 sleep such as spindles and SO are identified. The extracted features are sent to a machine learning predictor which outputs an estimation of the probability to be in N3. In parallel, the features are used together with time information to check if several hard conditions are met: do not stimulate before 15min after sleep onset; do not stimulate if a large movement happened less than 3 min ago; do not stimulate after 4h after sleep onset. If both the hard conditions are met and the N3 machine learning predictor outputs a probability to be in N3 larger than a threshold then 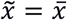, else 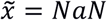 **E**- High-level block diagram of stimulation algorithm. First, the virtual channel is sent to a block, which estimates the phase of the signal in the delta band. We use a new phase detection algorithm here, which is described in the text body. Next the algorithm checks whether the phase is equal to 45°, the target that is set for stimulation, and emits a stimulation. However, there is a delay condition from the previous stimulation to ensure we do not stimulate at each SO. The two rules used here are: (i) do not stimulate more than 2 SO in a row, (ii) wait at least 9s before the previous pair of stimulations. If those delays conditions are met, then the stimulation order is given to the hardware, which emits 50ms stimulation through the bone conduction.

##### Quality gate

The quality gate allows the signal to proceed to the next stage if it reaches a threshold quality (Fig. 2B). This quality detector is a machine learning predictor (forest of decision trees) applied to a binary classification task on a large database of 2s windows labeled by sleep experts which specified if parts of the signal correspond to good or bad quality signal. Every 0.5s an estimation of the quality is made by this algorithm, returning a number between 0 and 1. If it crosses the minimal threshold then the signal is broadcasted to the channel switcher.

##### Channel switcher

The algorithm selects the channel with the highest quality (Fig. 2C). This so-called selected channel is referred as the ‘virtual channel’. A hysteresis switcher avoids switching to often from one channel to the other if they have similar quality.

##### N3 sleep gate

The N3 sleep gate classifies 30s windows of ‘virtual channel’ in N3 sleep *vs* else (Fig. 2D). This N3 sleep detector is mainly made of a machine learning predictor (forest of decision trees) fed with numerous features computed on the ‘virtual channel’ and on the accelerometer. For instance, we considered relative power in frequency bands on the EEG signal in intervals 0.4 to 4Hz, 4 to 8Hz, 8 to 12Hz and 12 to 18Hz, permutation entropy of EEG, and various measures of signal complexity. We also identify key sleep patterns in the signal such as spindles and slow oscillations. If the signal is detected as N3 sleep and meets hard conditions applied to avoid awaking the user, then it is broadcasted to the next stage. In other sleep stages, the data are not sent to the stimulation stage and no stimulation can be heard. Notably, the WDD does not stimulate if the quality of both channels is bad.

##### Phase fitting algorithm

The algorithm used here was inspired from the Cox et al. and consisted in fitting a sinus to the filtered 0.4 to 4Hz signal of the ‘virtual channel’ and identify the phase of the signal on the sinus itself [30]. In the considered case of a fixed frequency, this fit corresponded to a linear regression performed in real time and at each time step with a recursive least square method with a forgetting factor of λ= 0.99 providing a ‘memory’ (the equivalent of the size of a sliding window) of 5s. The fit was performed for 5 regularly spaced frequencies for the sinus between 0.8 to 1.2Hz, *i.e.* freq_list_= [0.8, 0.9, 1, 1.1, 1.2]. The frequency with the best fit was chosen. At each time step another sinus was fitted to the signal and only the last value was considered for stimulation. We used a Recursive Least Square method to perform the fitting economically at each time step [31] The frequency with the best fit was chosen. At each time step another sinus was fitted to the signal and only the last value was considered for stimulation. This resulted in a phase approximation, which is different than a normal sinus (Fig. 3). The algorithm in pseudo code with the following notations:

-t a column vector of regularly time steps between 0 and 2s at a sampling frequency with n=250 rows
- Λ is a diagonal matrix with diagonal (1, λ, λ^1^, λ^2,^… λ^(n-1).^).

**Fig. 3.**
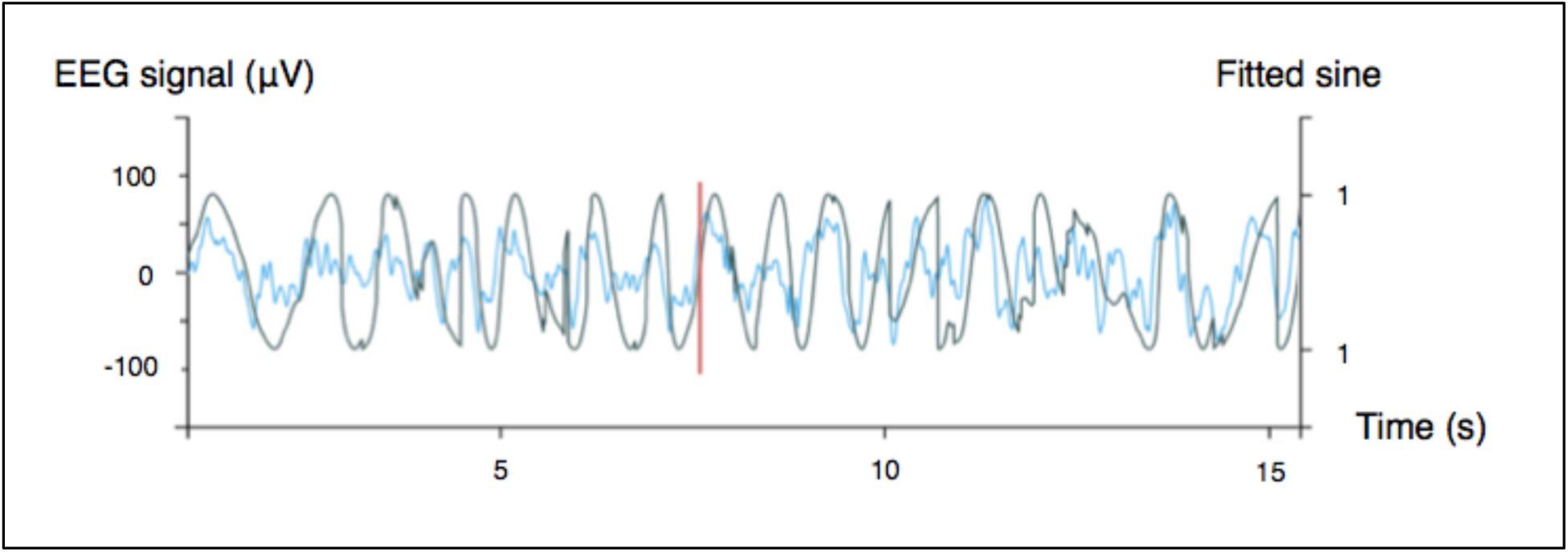
Illustration of the stimulation algorithm on a 15s epoch of EEG during N3 (blue). At each time, a sinus with the appropriate frequency is fitted to the last few seconds of the signal. Here, only the current value of the sinus is displayed (black) and serves as a basis for stimulation in the ascending phase. One stimulation trigger is shown in red.

Initialization

> for f in freq_list_:

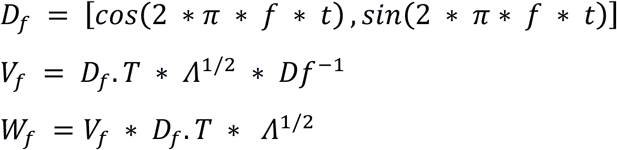

At each time step, for a new signal y

> for f in freq_list_:

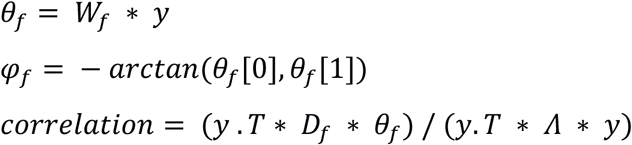

> Choose f and *φ*_*f*_ that maximize the correlation

This resulted in a phase approximation, which is different than a normal sinus (Fig. 3) since the estimated phase is recomputed at each time step.

##### Stimulation procedure

The stimulation was launched based on the estimated phase according to the previous procedure.

In part 1, the acoustic stimulation routine was inspired from previous studies and consisted in 2 consecutive SO phase-locked stimulations of 40 dB pink noise [16], [17]. The exact phase targeting was 45° in the ascending condition, right before the up-phase, since after some piloting, it seemed to be the optimal phase for driving SO.

Part 2 included only nights with stimulations (*i.e.* no nights with sham only) since the WDD’s users almost systematically turned on the stimulations when using the headband. However, in these so called “stimulation nights”, approximately 50% of the stimulations were real stimulations (*i.e.* with sounds set at 40 dB) and approximately 50% of the stimulations were sham (*ie* with no sound). Sham and real stimulations were randomly displayed through the night.

In both studies, a pause of 9s, minimum, between trains of 2 stimulations was made before detecting another SO and stimulating. Stimulations began after 15min of stable N3 sleep and persisted during this sleep stage solely, unless a movement or alpha rhythm was detected in the 6s following the stimulation. In which case, a 30s pause was initiated.

#### - A posteriori data analysis

##### Resynchronization procedure

In part 1, a resynchronization procedure was processed between the EEG signals provided by the PSG device and the WDD. It consisted in aligning the nights, but also to match their sampling frequency. Indeed, a slight change of sampling frequency in both devices occurred through the nights. These slight variations of less than 0.1Hz led to seconds of delay at the end of an 8h night. The variations in frequency changed non-linearly and non-monotonously during the night due to temperature changes. Thus, a sequential resynchronization procedure for chunks of 10min of recording was used where the problem was expressed as an optimization problem as a function of signal translation and sampling frequency.

##### Signals correlation methods

Correlation between PSG and the WDD was assessed on resynchronized signals with a Pearson correlation coefficient for windows of 2s. Signals with electrodes disbond were removed from the analysis (1.19% of the signal removed because of 2 bad PSG derivations, 4.72% because of 2 bad WDD derivations, 10.07% because of 1 bad PSG derivation and 13.11% because of 1 bad WDD derivation. The correlation between the PSG and the WDD could not be computed on the same derivations since the wiring of the WDD is unilateral (Fp1-M1, Fp2-M2). The classical wiring of the PSG was not changed into unilateral montage to avoid impairing the sleep stage classification of sleep experts who are used to a contro-lateral montage. Therefore, we compared the ‘virtual channel’ of both the WDD and the PSG. Eventually, the pair of channels that were compared always had a common location for one electrode, to the extent that both devices have to be set up to slightly different locations. Overall, this imperfect ‘virtual channel’ comparison underestimated the results of correlation and served as a lower bound to the real correlation.

##### Performance analysis of automatic N3 sleep detection

The performance analysis of the automatic N3 sleep detection of the WDD was assessed on the recordings from Part 1 by comparing the performance of the device to the manual sleep scoring of an expert on the PSG. The trained research technician was blinded to the conditions and scored the signals in accordance with AASM criteria [32] using SOMNOLOGICA (TM; Medcare, Reykjavik, Iceland). Stages annotations and timestamps were recorded. Note that none of the night analyzed here was involved in the training of the embedded automatic sleep staging algorithm. To assess the performance of the WDD, we determined the true positive rate *i.e*. correct N3 sleep detected by the WDD, false positive *i.e*. false N3 sleep detected, true negative *i.e*. correct N3 sleep rejected, false negative *i.e*. false N3 sleep rejected, sensitivity *i.e*. correct N3 sleep detection when the PSG also scores SWS) and specificity (*i.e*. ability of the WDD to measure false N3 correctly identified as such).

#### - Accuracy of the stimulations

The ability of the algorithm to target the positive half-wave (*i.e.* the ascending phase) of the SO was tested on the recordings from the clinical study (Part 1) to ensure that only stimulations elicited in N3 were analyzed. All the stimulations were summed up in a circular “polar plot” histogram by using a zero-phase digital filter with transfer function coefficients of a second order band-pass Butterworth filter in the delta band (0.4 to 4Hz). The phase angle at each pulse delivery was identified and a Hilbert transformed was applied on the EEG signal to identify the instantaneous phase at each pulse delivery. Circular histograms were created with 72 bins of 5° where 90° represents the peak of the upstate and the ascending targeted phase of stimulation delivery 45°.

#### - Event related potentials

The physiological impact of the stimulations on the EEG was assessed on the recordings from Part 2 in order to increase the statistical power by the important size of our sample (90 subjects, 10512 stimulations and 9872 sham triggers).

The averaged event related potentials (ERP) are presented as mean ± standard deviation In order to avoid phase delays, the spectral filtering of the signals was done with the same filters as described before but with a non-causal forward/backward scheme. The ERPs were time locked to both first and second trigger since as opposed to former algorithms; the duration between two stimulations could vary due to the particular shape of the signal. This method choice implies some non-causality in the filtered signals (*i.e.* may lead to significant difference prior the first stimulation) but guarantees no phase delay.

The impact of the stimulations over time was assessed by comparing (the stimulations of the 1st night – the sham of the 1^st^ night) to (the stimulations of the 10^th^ night – the sham of the 10^th^ night).

The power increase in the delta band was computed between stimulated and non-stimulated SO. More precisely, we computed the delta power in the 0.4 to 4Hz frequency band in a 4s window after the first stimulation (or sham) in each train of 2 stimulations (or shams). We used the squared norm of the discrete Fourier Transform of the 1024 time steps after the first trigger convolved with a Hann function. This provided 2 distributions of delta power: one after stimulations and one after sham. We then computed the percentage of increase between the mean of these two distributions.

When comparing different conditions, a T-test for the mean of the independent distributions corresponding to the two conditions was computed at each time step. This means that for each time bin distant from the first trigger or the second trigger, two distributions were collected: one for all the EEG values when there was stimulation, and another when there was no stimulation (sham). The time steps for which p<0.001 were considered as significant, meaning that the null hypothesis was rejected and that the distributions have a different mean.

## III. RESULTS

### A. Part 1-Validation of the acquisition and accuracy performances of the WDD in a clinical study

#### - Data characteristics

In part 1, data from four participants were discarded from the analysis: one due to defective headband, one due to damaged PSG, one due to poor sleep of the participant among the 4 nights which might be explained by the discomfort of the set-up and one due to the disrespect of the protocol regarding sleeping hours. This resulted of a final sample size of 20 subjects (7 women, mean age= 23.1 years, range 19-29 years, PSQI: 2.6 ±2.1, Beck: 1.2 ±2.0, HAD: 8.6 ±3.2) and 60 nights corresponding to the three conditions Sham, Ascending and Random conditions.

#### - Signal quality

Correlation scatters plot shows how two PSG channels correlate together and how the WDD correlates to PSG (Fig. 4). The Pearson correlation between WDD and PSG signal show a maximum around the value of 0.6.

**Fig. 4.**
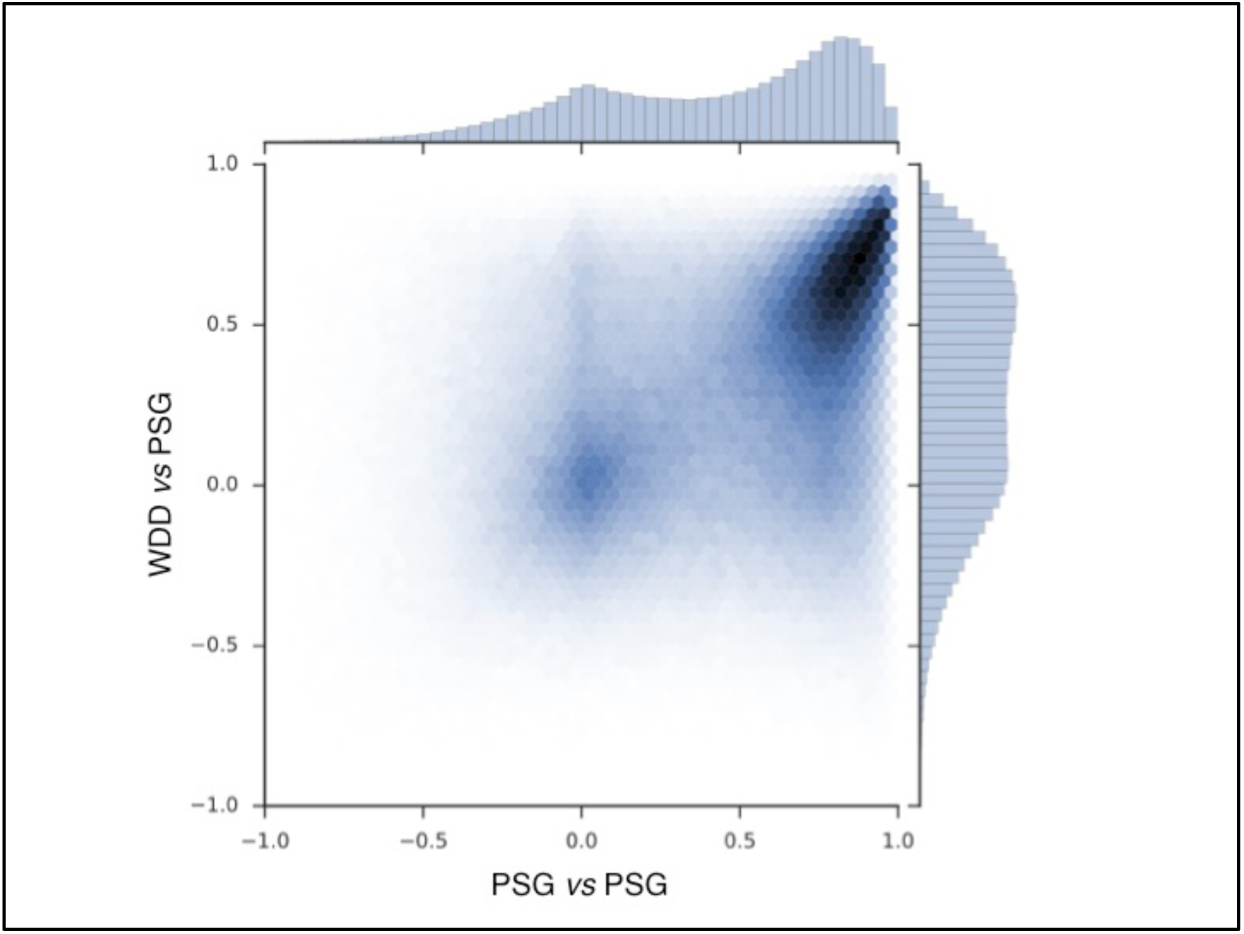
Pearson correlation scatter plot for 697017 windows of 2s with resynchronized PSG and the WDD recordings. PSG vs PSG shows the correlation between the two frontal channels of the PSG device. PSG vs the WDD shows the correlation between the virtual channels of the WDD and PSG channel.

Illustrative samples of the signal obtained with the PSG and the WDD for each sleep stage are presented in Fig. 5. The typical rhythms including alpha and theta as well as the typical sleep patterns such as spindles, K-complexes and SO were distinguishable in both recordings.

**Fig. 5.**
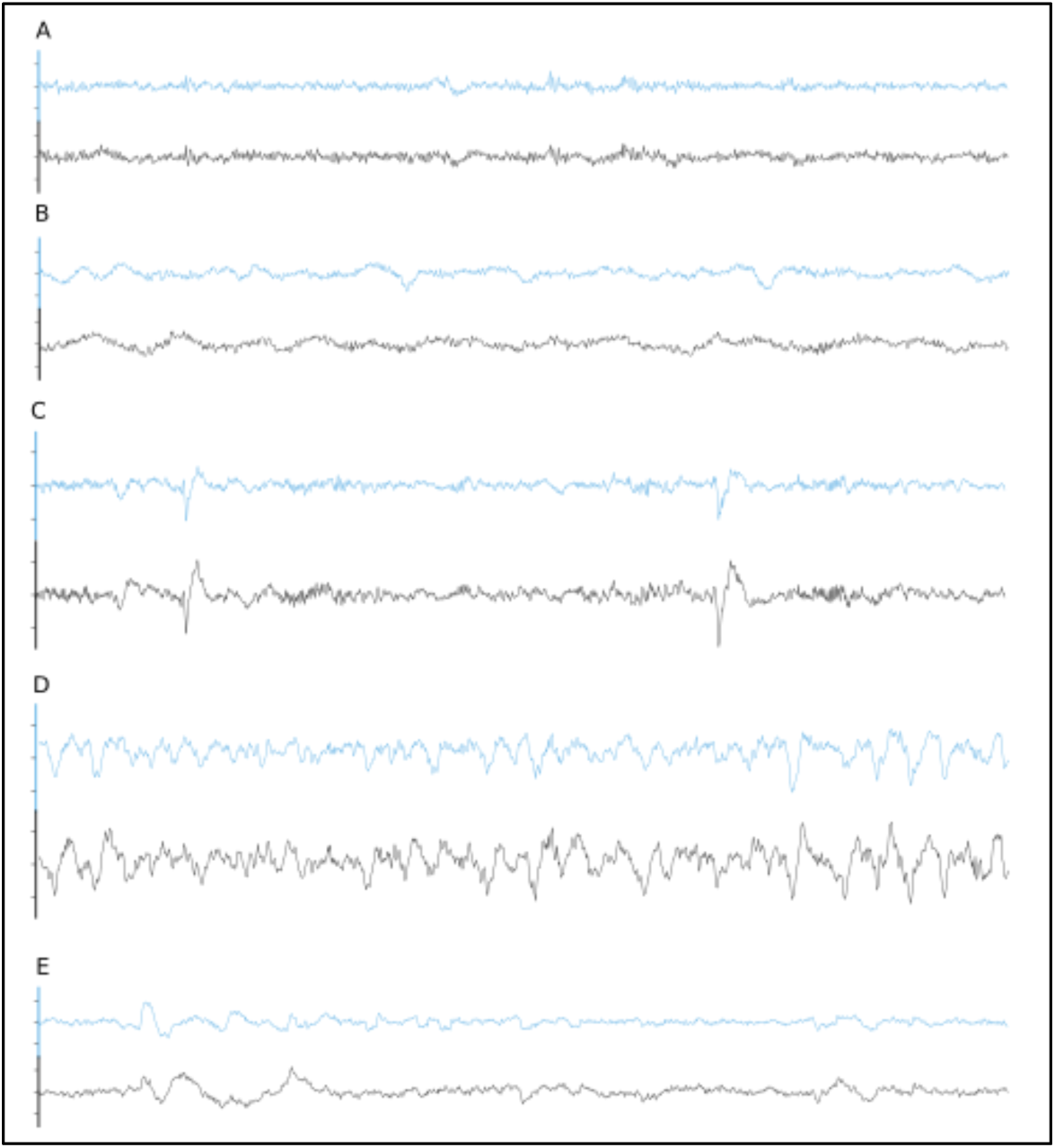
Representative 30s epoch of A- Wakefulness, B- N1, C- N2, 3- N3, 4- REM obtained with the simultaneous recording of the WDD (blue) and the PSG (black).

Time-frequency plots show very similar distribution of frequencies across the night when comparing the two devices (Fig. 6 for a representative plot, see all the individual plots in Annex 1).

**Fig/ 6.**
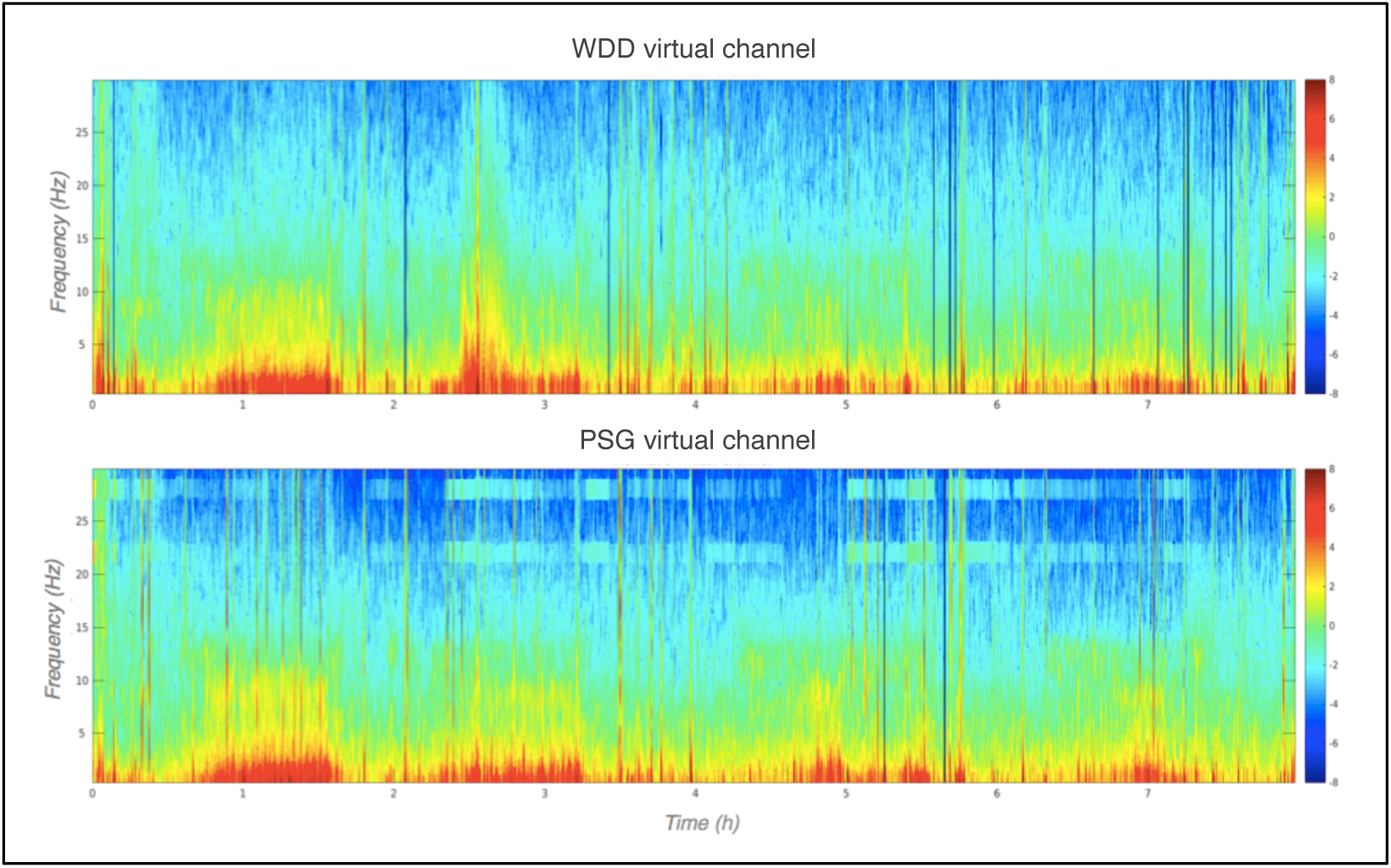
Representative multitaper EEG spectrogram of a full sleep night from the WDD (on top) and on the PSG (at the bottom) recordings.

#### - Automatic N3 detection

The performance of WDD to detect automatically N3 sleep with an algorithm as compared to the traditional sleep staging provided by the sleep expert on the PSG show high specificity (0.90), compared to sensitivity (0.70). Out of 42302 total epochs scored, 12276 epochs were scored in N3 with 3017 epochs appearing as false positive, 3666 as false negative, 8610 as true positive and 27009 as true negative (Table I).

**TABLE I:**
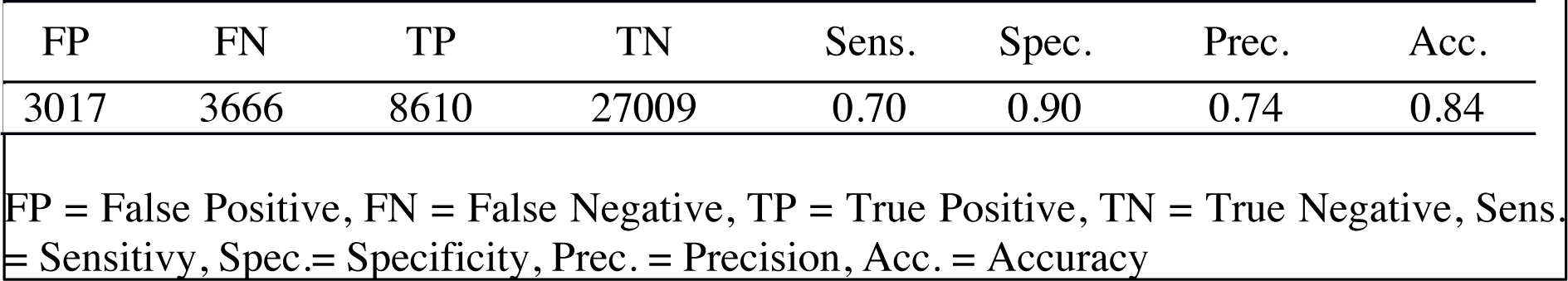
Performance of The WDD’s Automatic Sleep Stage Algorithm

| TABLE I<br>PERFORMANCE OF THE WDD'S AUTOMATIC SLEEP STAGE ALGORITHM |  |  |  |  |  |  |  |
| --- | --- | --- | --- | --- | --- | --- | --- |
| FP | FN | TP | TN | Sens. | Spec. | Prec. | Acc. |
| 3017 | 3666 | 8610 | 27009 | 0.70 | 0.90 | 0.74 | 0.84 |
| FP = False Positive, FN = False Negative, TP = True Positive, TN = True Negative, Sens. = Sensitivity, Spec.= Specificity, Prec. = Precision, Acc. = Accuracy |  |  |  |  |  |  |  |

A total of 17579 stimulations were elicited by the device and 17786 Sham. We observed that 86.1 % of stimulations or sham were elicited in N3, 11.0 % in N2, 0.4 % in N1 1.4 % in REM and 1.1 % during wakefulness according to the sleep expert (Table 2. See Annex 2 for the individual hypnograms and the stimulations triggers).

**TABLE II:**
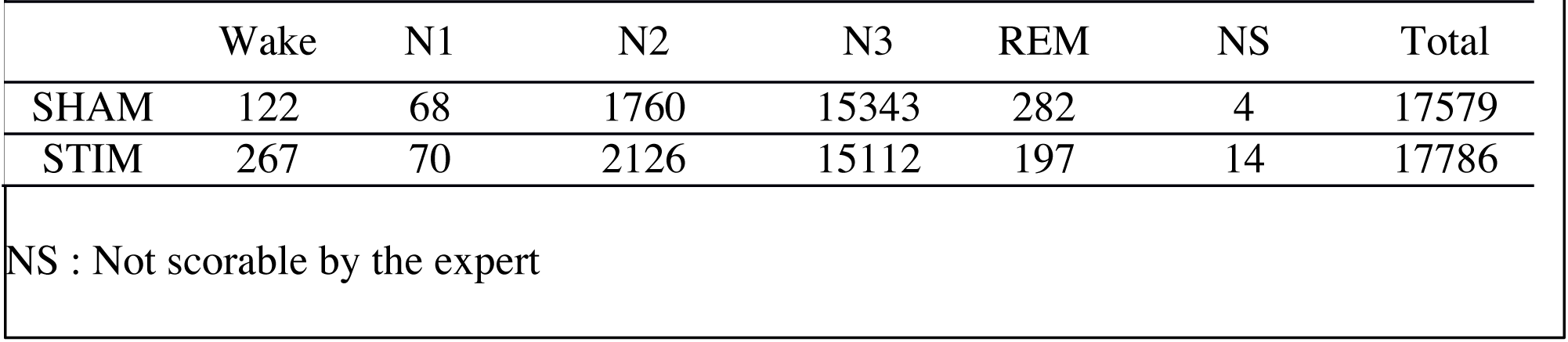
Number of stimulations by sleep stage according to the sleep expert scoring

|  |  |  |  |  |  |  |  |
| --- | --- | --- | --- | --- | --- | --- | --- |
|  | Wake | N1 | N2 | N3 | REM | NS | Total |
| SHAM | 122 | 68 | 1760 | 15343 | 282 | 4 | 17579 |
| STIM | 267 | 70 | 2126 | 15112 | 197 | 14 | 17786 |

NS : Not scorable by the expert

#### - Stimulation accuracy

The average time of stimulation for the 7059 pink noise at the ascending phase was 45±52° (Fig. 7) (Fig. 7, see Annex 3 for individual polar plot).

**Fig. 7.**
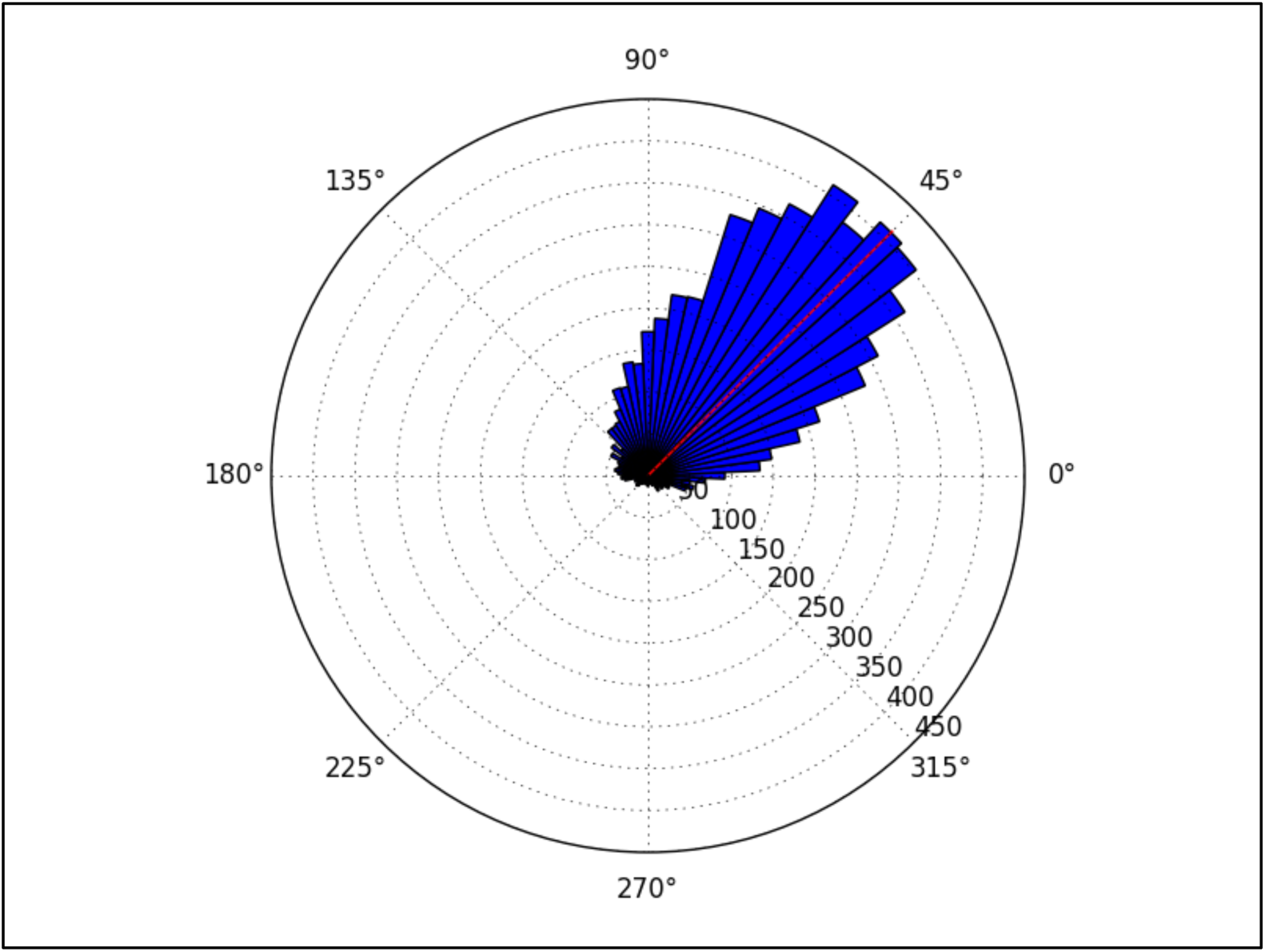
Polar histogram showing 7059 stimulations as a function of the phase of the signal. The targeted phase was 45° which represents the middle of the ascending slope. 90 degrees corresponds to the peak of the up state, 270 degrees to the trough of the down state.

### B. Part 2 - Electrophysiological effects of auditory closed loop stimulation provided by the WDD in an observational pilot study

#### - Data characteristics

In part 2, after applying the selection criteria to ensure the quality of the nights (see material and methods), 90 subjects (9 women, mean age: 40.8±10.9 years old) were included for ERP impact of auditory stimulations. The longitudinal analysis involving 10 nights of stimulations in a raw only included 20 subjects (1 woman, mean age: 45.4±8.0 years old). Indeed, since this study was observational, subjects were not asked to wear repetitively the WDD and most of them wore it sparsely (couple of days a week).

#### - Electro-physiological impact of the auditory stimulations

Across all nights and all stimulations a total of 10512 stimulations and 9872 sham triggers were displayed, The averaged ERP time-locked to the first (Fig. 8A) and to the second stimulations (Fig. 8B), elicited a greater increase in the amplitude of slow oscillatory activity in the Stim condition, as compared to sham, with this effect tapering off after the second oscillation (p<0.001).

**Fig. 8.**
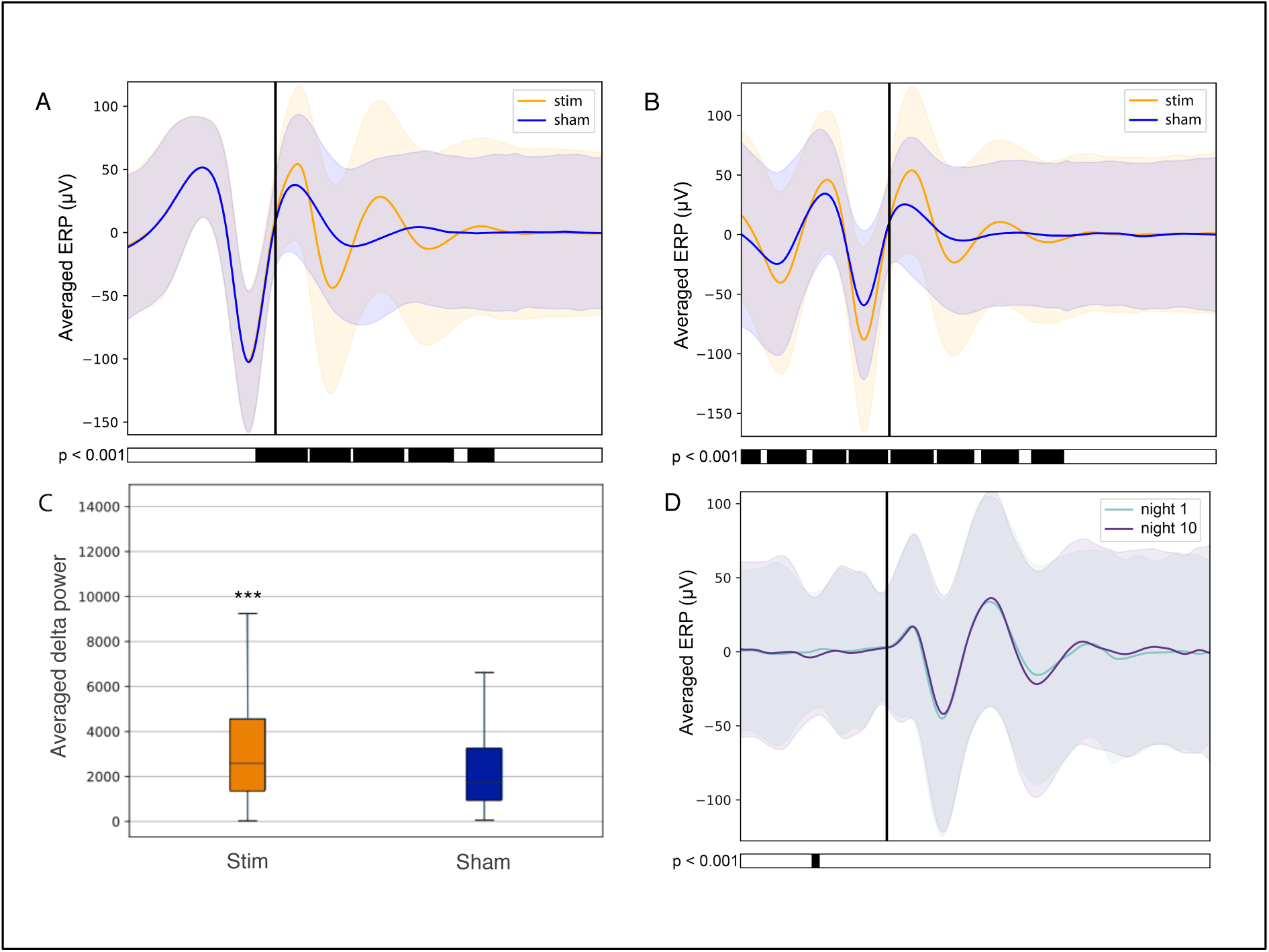
**A, B-** Averaged ERP (±SD) time locked to the first (**A)** and second **(B)** stimulus for the stim (orange line) and sham (blue line) in the observational study (Study 2). **C-** Averaged power in the delta band in the 4s following the 1^st^ stimulation (Stim) or sham trigger (Sham). **D-** Resulting ERP of the 1^st^ (light blue line) and 10^th^ night (purple line) stimuli where the“1^st^ night” and the “10^th^ night” refer to the difference between the stim and the sham of the first and tenth night respectively. Black line indicates the stimuli trigger. Black bars indicate time points where differences between the two conditions were statistically significant (p<0.001). *** indicates significant difference between the Stim and the Sham condition (p<0.001).

On average, the power increase in the delta band in the 4s following the first stimulation was of 39.47 % when stimulating as compared to sham triggers (Fig. 8C).

Finally, no difference was observed on averaged ERP after wearing the device for 10 nights in a raw - *ie* when comparing the impact of the 1st night of stimulations to the 10^th^ night with stimulation (Fig. 8D).

## IV. DISCUSSION

The present research aimed at assessing the performances of the WDD, an ambulatory dry-electrodes EEG device, for auditory closed-loop stimulations of SO during N3 sleep in the home environment. Here, we reported its technical performance from a clinical trial including 20 healthy participants. In this first part, we showed that the device had a good acquisition quality compared to a PSG with a good ability to detect N3 sleep in real time (specificity: 0.90, sensibility: 0.70) and a precise algorithm for auditory-closed loop stimulations (45±52° reached on average for a 45° targeting). Then, we observed that auditory closed loop stimulations on a large number of participants in the home environment led to similar results than anterior research on small samples. Finally, we addressed for the first time the impact of N3 stimulations over 10 nights and showed that the EEG responses to the stimulations after 10 nights were not different from the ones in the first stimulation night.

The experimental procedure followed during part 1 led to good results in terms of acquisition with good visual identification of sleep patterns (Fig. 5), good correlation between signals when using Pearson correlations in 2s windows (Fig. 4) and when comparing the all night spectrogram (Fig. 6).

While in the clinical practice, considerable time and large attention are deployed for the EEG set-up, it is encouraging to observe that subjects were able to use the WDD by themselves to both launch the recording and place the headband. The autonomous placement of the headband by the subjects might have lead to a small offset compared to the optimal position of the electrodes for the comparison between the two devices. This displacement might have contributed to the values dispersion on the Pearson correlation scatter plot (Fig. 4).

The detection of N3 sleep, as compared to a PSG gold-standard, led to good results with a specificity of 0.90 and a sensitivity of 0.70 (Table I), which has to be put in perspective with the fact that the inter-scorer variability for sleep stage classification along the AASM rules is about 82% [33]. These results were obtained with dry frontal electrodes referred to mastoids whereas the PSG encompassed EEG, EOG and EMG. Indeed, post-processed automatic sleep staging using EEG, EOG and EMG reported performance of 0.92 for specificity and 0.74 for sensitivity [34]. A similar commercial device, the Zeo Wireless using frontal electrodes solely, showed a specificity of 0.62 [35] as compared to Somnolyzer, a standard automatic sleep stager [36]. Importantly and as opposed to most results in the literature, the WDD algorithms described here run in real-time and are fully embedded on the headband with no outside communication (*e.g.* Bluetooth, Wi-Fi…) during the night. This imposed significant optimization constraints on all computations performed and oriented drastically the nature of the algorithms used: forest of decision trees rather than deep learning approach. Nonetheless, good performances were reached mainly due to the vertical technological integration of the WDD, which led to a precise optimization of the algorithm.

Auditory closed-loop studies mostly agree that the timing of SO stimulations matters [16], [18], [37]. Therefore, studies were led to improve the stimulations algorithm. In our study, we aimed at delivering auditory stimulations in the ascending phase *i.e.* 45° of the SO. On average, our algorithm reached 45±52° (Fig. 7). To our knowledge, this performance is higher than the phase locked loop (PLL) algorithms previously published. In the initial auditory closed-loop study, the up phase of the stimulation was targeted but the data to assess the precision of the algorithm were not available [15], [16]. In 2014, the PLL algorithm used in the study of Cox and collaborators aimed to target the SO’s up and down phases [18]. The performance for the up phase (90°) targeting reached 79±66° on average which makes a 11° difference from the targeted placement. More recently, a study showed that the stimulation accuracy was improved by stimulating at 50±27° for a targeted placement of 60° in a protocol including sleep naps [17]. Another recent study achieved a very good performance in elderly participants and reached 329±73° on average for a 340° targeting [20]. Notably, all these studies were led in a sleep lab with an experimental setting requiring the use of a complex wiring connected to a computer and sometimes even sleep technicians to initiate the stimulation algorithm when N3 occurred. To our knowledge, only one study used a possible ambulatory set-up where computations were processed on a tablet right next to the bed [21]. The performance of the algorithm in this study showed a difference of 18±67° from the desired phase.

The good acquisition performance made possible the ability to stimulate during N3 sleep precisely on the ascending phase of the SO on a large number of participants (90 participants in Part 2). As observed in previous studies including about 10 to 20 participants [15], [16], [20], [21], [38], the auditory closed-loop stimulation inspired by Ngo’s protocol over our 1000 nights led to an increase in amplitude during the period immediately following the stimulation. In particular, the power increase of the signal in the delta band (0.4 to 4Hz) increased 4s after a stimulation suggesting a strong local impact. As displayed in Fig. 8, the averaged ERPs showed a strong amplification due to stimulations. This is to be understood as a synchronization of brain SO on the stimulations showing a strong local interaction between stimuli and brain activity even during N3 sleep. The increased standard deviation due to stimulations displayed in Fig. 8 showed an increase in the amplitude of the SO which gets back to its normal states after ∼5s. The analysis of the longitudinal effects of auditory closed-loop stimulations through 10 consecutive nights showed no significant difference as compared to the stimulation of a single night, suggesting that no adaptation mechanism occurs to regulate the impact of the stimulation through the night or with daily stimulations. These results suggest that the auditory stimulations provided by the bone conduction instead of habitual headphones or loudspeakers were similarly able to activate the non-lemniscal pathway and thus trigger slow waves in response to the auditory stimulus [10]. This finding is rather encouraging since it implies that co-sleepers could be individually stimulated with the WDD, which would not have been possible with traditional montage.

In this observational study, data were issued from a «conservative» approach method consisting in choosing stimulations parameters to avoid awaking up the WDD’s users as a first objective. Therefore the volume was kept low (40dB), the number of stimulations was moderate during a night (∼100 per nights) and stimulations were exclusively triggered during N3 sleep. Moreover, we deliberately tuned our N3 detection algorithms to reach a high specificity at the expense of sensitivity. Indeed, to limit potential awakenings due to sound stimulation in other sleep stages, the WDD was designed to be particularly good at detecting periods identified as being different from N3 sleep. In other words, it is crucial that the algorithm does not make mistakes at declaring that a given period is N3 sleep, even at the expense of missing some ambiguous periods. As discussed by Bellesi et al., we believe that the optimization of stimulations parameters such as the targeted sleep stage (N2), the volume intensity, the type of sound, the phase of the stimulation and the number of stimulations (overall or in a single train) could possibly lead to better deep sleep enhancement [1]. Moreover, the age of the subject might also change the way the brain respond to auditory stimulation since the brain and thus the sleep EEG is already different. A personalization approach may thus maximize the effect of auditory stimulations and lead to different results as the one we presented after a single or several stimulation nights.

As a limit, the observational study suffers from the absence of control over the subject’s behavior. Subjects bought the WDD and used it on voluntarily basis. There was no proper recruitment or screening and we had little information about their profiles and habits. In particular, their sleep habits, disease, drugs consumption *etc*…were not verified. Also it has to be noted that a recruitment bias is possible since subjects who keep using the WDD after numerous nights may also be the subjects whose sleep was responsive to the stimulations. On the contrary, we had fewer recordings from users with a low sensitivity threshold whose sleep may be negatively altered by the stimulations. These constraints were inherent to any observational ecological study, and may alter some findings about the global population of users. However, since we restricted our analysis to the quantification local physiological impacts of the stimulations, we do not believe that these biases significantly impacted the results presented here. Nonetheless a systematic study covering the longitudinal effects of stimulation as well as studies including more women is needed to confirm our results.

## V. CONCLUSION

This study showed that the WDD - a fully integrated dry-electrode commercial wearable could monitor sleep on frontal EEG derivations with a good acquisition quality compared to gold standard PSG devices. The specificity and sensibility to detect N3 sleep as well as its stimulation accuracy were above the performances found in the literature. Its use in the home environment has resulted in an unprecedented number of nights. We were able to replicate anterior studies on EEG response to auditory closed-loop stimulations and showed for the first time that these stimulations over 10 nights did not reduce nor potentiate the EEG responses of a single stimulation night.

Overall, given its performance and its ease of use, the WDD may be an excellent way to go further into the analysis of N3 sleep stimulations, including targeting memory reactivation, on larger population than in anterior works. More broadly, it provides new perspectives on the assessment of sleep EEG biomarkers by possibly being a substitute of PSG in “outside of the lab” longitudinal cohort studies.

## ACKNOWLEDGMENT

We would like to thank the Sleep and Fatigue Team including Bougard C, Dorey R, Drogou C, Drogou G, Erblang M, Gomez-Merino D, Rabat A and Van Beers P as well as Ferret M, Voluntario V and Dr Giordanella for their help and commitments in this study. We also would like to thank the subjects who take part in the study and all the Rythm team for their commitments in working on the headband.

